# Structure of a G protein-coupled receptor with GRK2 and a biased ligand

**DOI:** 10.1101/2022.10.19.512855

**Authors:** Jia Duan, Heng Liu, Yujie Ji, Qingning Yuan, Xinzhu Li, Kai Wu, Tianyu Gao, Shengnan Zhu, Wanchao Yin, Yi Jiang, H. Eric Xu

## Abstract

Phosphorylation of G protein-coupled receptors (GPCR) by GPCR kinases (GRKs) desensitizes G protein signaling and promotes arrestin signaling, which is also modulated by biased ligands^1–6^. Molecular assembly of GRKs to GPCRs and the basis of GRK-mediated biased signaling remain largely unknown due to the weak GPCR-GRK interactions. Here we report the complex structure of neurotensin receptor 1 (NTSR1) bound to GRK2, Gαq, and an arrestin-biased ligand, SBI-553^7^, at a resolution of 2.92 Å. The high-quality density map reveals the clear arrangement of the intact GRK2 with the receptor, with the N-terminal helix of GRK2 docking into the open cytoplasmic pocket formed by the outward movement of the receptor TM6, analogous of the binding of G protein to the receptor. Strikingly, the arrestin-biased ligand is found at the interface between GRK2 and NTSR1 to enhance GRK2 binding. The binding mode of the biased ligand is compatible with arrestin binding but is clashed with the binding of a G protein, thus provide an unambiguous mechanism for its arrestin-biased signaling capability. Together, our structure provides a solid model for understanding the details of GPCR-GRK interactions and biased signaling.

---

GPCRs comprise the largest family of cell surface receptors whose signaling is primarily mediated by two types of downstream effectors: G-proteins and arrestins. The switch of GPCR signaling from G-protein pathways to arrestin pathways is controlled by a small family of GPCR kinases, GRKs, which phosphorylate either the receptor C-terminal tail or the third intracellular loop (ICL3)^1–3^. Phosphorylation of GPCRs promotes recruitment of arrestin, which blocks G-protein binding and desensitizes G-protein signaling^3^. Because drugs that selectively activate either G-protein pathways or arrestin pathways (biased signaling) are proposed to have better therapeutic and safety index^4,5^, the mechanism of GPCR biased signaling has been a subject of intensive research over the past two decades.

GRK2, along with GRK1, are the prototypes of GRKs that belong to the AGC family of serine/threonine kinases^8–10^. There are seven GRKs, which can be grouped into the rhodopsin kinase subfamily (GRK1 and GRK7), the β-adrenergic receptor kinase subfamily (GRK2 and GRK3), and the GRK4 subfamily (GRK4, GRK5 and GRK6)^9^. All GRKs share conserved sequence features and structural arrangements^11^. At the N-terminus is a conserved segment that formed a helix in the active GRK structures, followed by the first eight helices of a regulatory G-protein signaling homology domain (RHD)^10,12^. The kinase domain (KD) is inserted into a loop between helices 8 and 9 of RHD, a conserved domain of a nine-helix bundle found in regulatory G-protein signaling proteins^13^. Following the kinase domain and helix 9 of RHD are the less conserved C-terminal GRK domains, which are mainly responsible for membrane binding^14^. In the case of GRK2, its C-terminus contains a pleckstrin homology domain (PHD) that interacts with Gβγ subunits of G protein^15^. The RHD of GRK2 also interacts with Gαq when it is in complex with GTP^16^. The binding of both Gαq and Gβγ subunits to GRK2 facilitates its membrane association^15,16^.

As GPCR signal transducers like G proteins and arrestins, GRKs are rest at the basal state, and can be recruited and activated by active GPCRs^3^. The molecular basis of how GPCR signal transducers recognize and regulate GPCR signaling has been a research focus of GPCR structural biology^14,17–21^. Structures of many GPCR-G protein complexes and GPCR-arrestin complexes have been solved, which reveal that both G proteins and arrestins recognize the open cytoplasmic pocket induced by the outward movement of TM6 in the activated GPCRs^19,20,22–25^. Due to much weaker interactions between GPCR and GRKs, high resolution structure of a GPCR-GRK complex is technically challenging. A structure of rhodopsin in complex with GRK1 has provided a breakthrough view of the overall assembly of GRK1 with rhodopsin via its N-terminal helix^18^. However, the relatively low resolution of the structure lacks the density for the conserved RHD domain of GRK1 and limits the detailed understanding of rhodopsin-GRK1 interactions and GRK1 activation by the active rhodopsin.

Neurotensin receptor 1 (NTSR1) is a class A GPCR that is regulated by an endogenous peptide ligand, neurotensin (NTS)^26^. Up on activation, NTSR1 couples to various signal effectors, including several subtypes of G proteins, GRKs, and arrestins, to mediate neurotransmission and neuromodulation in the central nervous system^26–28^. Because of its diverse physiological roles, NTSR1 has been proposed as a drug target for addiction, obesity, analgesia, cancer, Parkinson’s disease, and schizophrenia^7^. Structures of NTSR1 in complex with Gi or beta-arrestin have been determined by cryo-electron microscopy (cryo-EM)^22,23,29^. Notably, SBI-553, a β-arrestin-biased allosteric ligand of NTSR1 that antagonizes G-protein signaling, selectively reduces addictive behaviors without the unwanted side effects of hypotension, hypothermia, and motor impairment, which are typically associated with balanced agonism of NTSR1 induced by neurotensin^7^. However, the structural basis of β-arrestin-biased agonism of SBI-553 remains unknown. In this paper, we report the structure of NTSR1 bound to NTS, GRK2, Gαq, and SBI-553 at a resolution of 2.92 Å, which reveals detailed interactions between NTSR1 and GRK2, and provides a molecular explanation for the β-arrestin-biased agonism of SBI-553.

## Complex assembly and structure determination

To identify a stable GPCR-GRK2 complex, we used Tango assays^30^ to screen various members of class A GPCRs, and NTSR1 turned out to be one of the strongest receptors that interact with GRK2 (Extended Data Fig. 1a). Addition of SBI-553 further increased NTSR1-GRK2 interaction (Fig. 1a). The presence of SBI-553 enhanced potency and efficacy of NTS to promote GRK2 recruitment to NTSR1 (Fig. 1b). Coexpression of NTSR1 with GRK2 as well as Gαq and Gβγ formed a complex that could be purified to homogeneity but it was unstable (Extended Data Fig. 1b). We introduced the NanoBiT tethering strategy^31,32^ to stabilize the complex by fusing LgBiT to the C-terminus of NTSR1 and HiBiT to the C-terminus of GRK2. The purification of the above complex showed a sub-stoichiometry ratio of the Gβγ subunit (Extended Data Fig. 1c), indicating instable association of the Gβγ subunit with the rest of the complex. We thus omitted the Gβγ subunit from the final complex assembly, which was further stabilized by chemical crosslinking with BS_3_ for cryo-EM studies (Extended Data Fig. 1d-f).

**Fig. 1:**
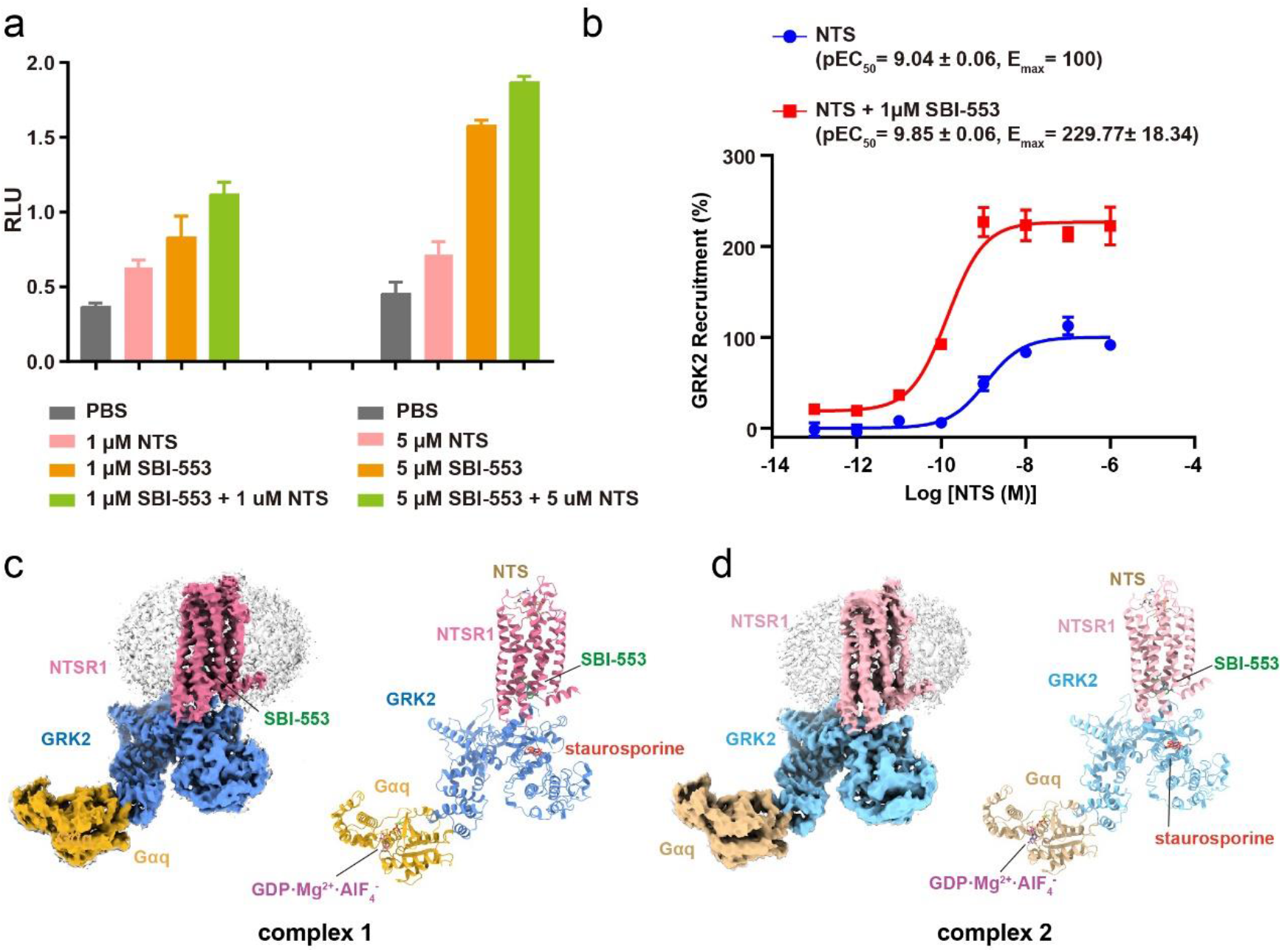
Cryo-EM structures of the NTSR1-GRK2-Gαq complexes. **a**, NTS and SBI-553 improve NTSR1-GRK2 interaction determined by Tango assay. RLU, relative luciferase units. **b,** NTSR1-GRK2 recruitment promoted by addition of NTS and SBI-553 determined by NanoBiT assay. Data were shown as mean ± S.E.M. from three independent experiments (n=3), performed in triplicates. The representative concentration-response curves were shown. **c, d,** Cryo-EM density maps and ribbon presentation of the NTSR1-GRK2-Gαq complexes. Complex 1 (**c**) and complex 2 (**d**).

A total of 57,477 film images were collected, which yield ~40 million initial particles. Further 2D and 3D classifications generate two maps at resolutions of 2.92 Å and 3.09 Å (Extended Data Fig. 2). The data and structure statistics are summarized in Extended Data Table 1. Both maps were sufficiently clear to place NTSR1, NTS, GRK2, Gαq, and the bound SBI-553, staurosporine, and GDP·AlF4^-^·Mg^2+^ (Fig. 1c, 1d and Extended Data Fig. 2, 3). Comparison of these two complexes reveals that they have very similar NTSR1 structure but a swing of GRK2 of ~5-6 Å related to NTSR1 (Extended Data Fig. 4), suggesting the dynamics of the NTSR1-GRK2 complex assembly. 3D variability analysis (3DVA) of the two cryo-EM maps also reveal dynamic swing of GRK2 around NTSR1, especially the Gαq subunit and the relative positions between the RHD and kinase domain (Extended Data movie). Complex 1 (Figure 1c) has higher resolution and is thus used for detailed analysis below.

## Structure of NTSR1-GRK2-SBI-553 complex

Within the complex structure, NTSR1 resembles the NTSR1 structure in complex with Gi and β-arrestin^22,23,29^ (Fig. 2a), with an overall RMSD less than 1.0 Å for the entire Cα atoms of NTSR1. Compared to the inactive NTSR1 structure, conformational changes mainly occurred at cytoplasmic ends of TM5 (4.5 Å shift), TM6 (11.3 Å shift) and TM7 (1.7 Å shift), consistent with an active conformation of NTSR1 (Fig. 2a).

**Fig. 2:**
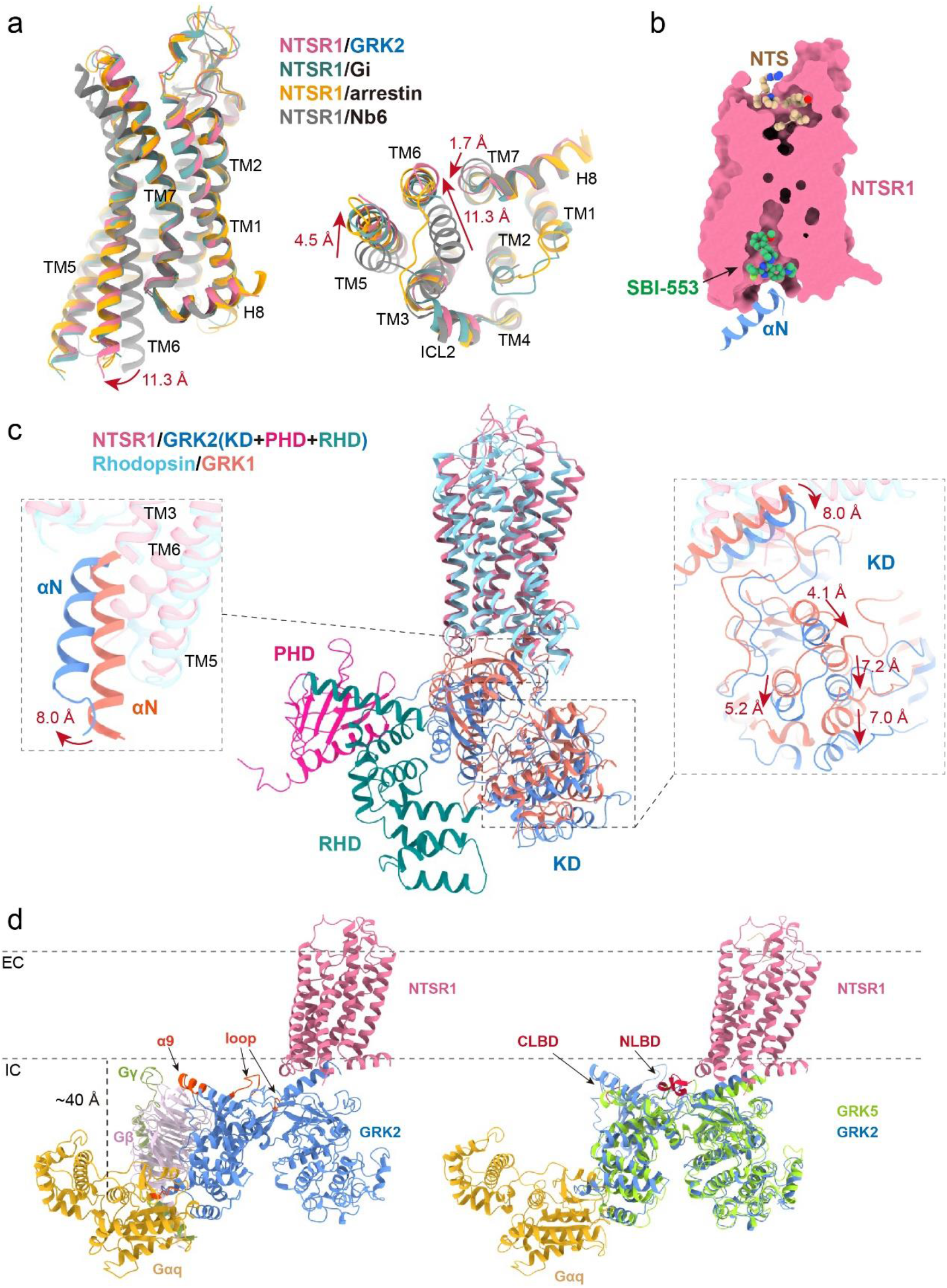
Structural features of the NTSR1-GRK2-Gαq complex. **a,** Structural comparison of the NTSR1 from NTSR1-GRK2-Gαq complex with the inactive NTSR1 (PDB code: 7UL2), NTSR1 from NTSR1-arrestin2 complex (PDB code: 6UP7) and NTSR1 from NTSR1-Gi complex (PDB code: 6OS9). **b**, The overall arrangement of the NTS and SBI-553 binding pockets in NTSR1. **c**, Structural comparison of the NTSR1-GRK2-Gαq complex with the rhodopsin-GRK1 complex (PDB code: 7MTA). KD, kinase domain. PHD, pleckstrin homology domain. RHD, regulatory G-protein signaling homology domain. **d,** Possible membrane binding sites of GRK2 and GRK5. The possible lipid binding sites from GRK2 are highlighted in orange, and the N-terminal lipid binding site (NLBD), the C-terminal lipid binding site (CLBD) of GRK5 (PDB code: 6PJX) are shown in dark red. α9, helix 9. EC, extracellular membrane layer. IC, intracellular membrane layer. The distance between the N-terminal of Gαq with the intracellular membrane layer is around 40 Å, which could be reached by the stretch loop of 38 residues from the N-terminus of Gαq.

At the extracellular side, NTS, the peptide ligand, is fit into the top-central TMD pocket (Fig. 2b). At the intracellular side, SBI-553 is found at the bottom-central cytoplasmic pocket, Underneath SBI-553 is the N-terminal helix of GRK2, which docks into the open cytoplasmic pocket. The overall structure of NTSR1-GRK2 complex is similar to the rhodopsin-GRK1 complex^18^ (Fig. 2c), however, the position of the N-terminal helix of GRK2 is shift by as much as 8.0 Å relative to the N-terminal helix of GRK1 (Fig. 2c). Correspondingly, the whole kinase domain of GRK2 is shift by as much as 7-8 Å from the GRK1 kinase domain (Fig. 2c).

Compared to the partial GRK1 structure in the rhodopsin-GRK1 complex, GRK2 from the NTSR1-GRK2 complex has nearly complete structure with RH and PH domains clearly defined in the structure (Fig. 2c). In the structure, Gαq is bound to the RHD of GRK2 (Fig. 2d). Comparing the GRK2 structure from the NTSR1 complex to the crystal structure of GRK2 from the complex with Gαq and Gβγ reveals three major differences as below^16^ (Extended Data Fig. 5). The GRK2 structure from the NTSR1 complex contains a N-terminal helix that is packed onto the kinase domain (Extended Data Fig. 5), has a breakage in the ionic lock between its RHD from the KD, and adopts a closed conformation in its kinase domain that is in the active state (Extended Data Fig. 5). In contrast, the GRK2 crystal structure from the complex with Gαq and Gβγ does not have the N-terminal helix, contains the ionic lock between its RHD and the KD as seen in the GRK5 structure^14,33^, and adopts an open conformation in its kinase domain, resembling the inactive state (Extended Data Fig. 5).

The overall arrangement of the NTSR1-GRK2 complex also present possible association of GRK2 with the membrane lipid layer (Fig. 2d). Alignment of H8 of NTSR1 with the membrane layer reveals that the C-terminal tip of helix 9 from RHD, the loop between β-strands 1 and 2, and the loop between β-strands 5 and 6 from PHD are in close contact with the membrane layer (Fig. 2d, Extended Data Fig. 6). Additional binding of GRK2 to the membrane layer could come from lipid modifications in the Gα and Gγ subunits (Fig. 2d). Modeling of the Gβγ subunit into the NTSR1-GRK2 structure suggest that the C-terminal lipid modification of the Gγ subunit is also close to the membrane layer (Fig. 2d). In addition, superposition of GRK5 to GRK2 in the NTSR1-GRK2 complex reveals that the N-terminal lipid binding domain (NLBD) and C-terminal lipid binding domain (CLBD) of GRK5 are near the membrane layer (Fig. 2d, Extended Data Fig. 6), consistent with their roles in lipid binding^14^.

## The GRK2-NTSR1 interface

The GRK2-NTSR1 interface is at the center of the complex, which has relatively high resolution at ~2.5 Å (Extended Data Fig. 2c, 2d), thus the density map is clear for interface residues, which reveals detailed intermolecular interactions between GRK2 and NTSR1 at the residue-specific levels (Fig. 3). The GRK2-NTSR1 complex has one major interface comprised by the N-terminal helix of GRK2, which inserts into the open TM6 pocket (Fig. 3a, 3b), and one minor interface comprised by ICL2 of NTSR1 that interact with the loop between the N-terminal helix and the RHD (Fig. 3c).

**Fig. 3:**
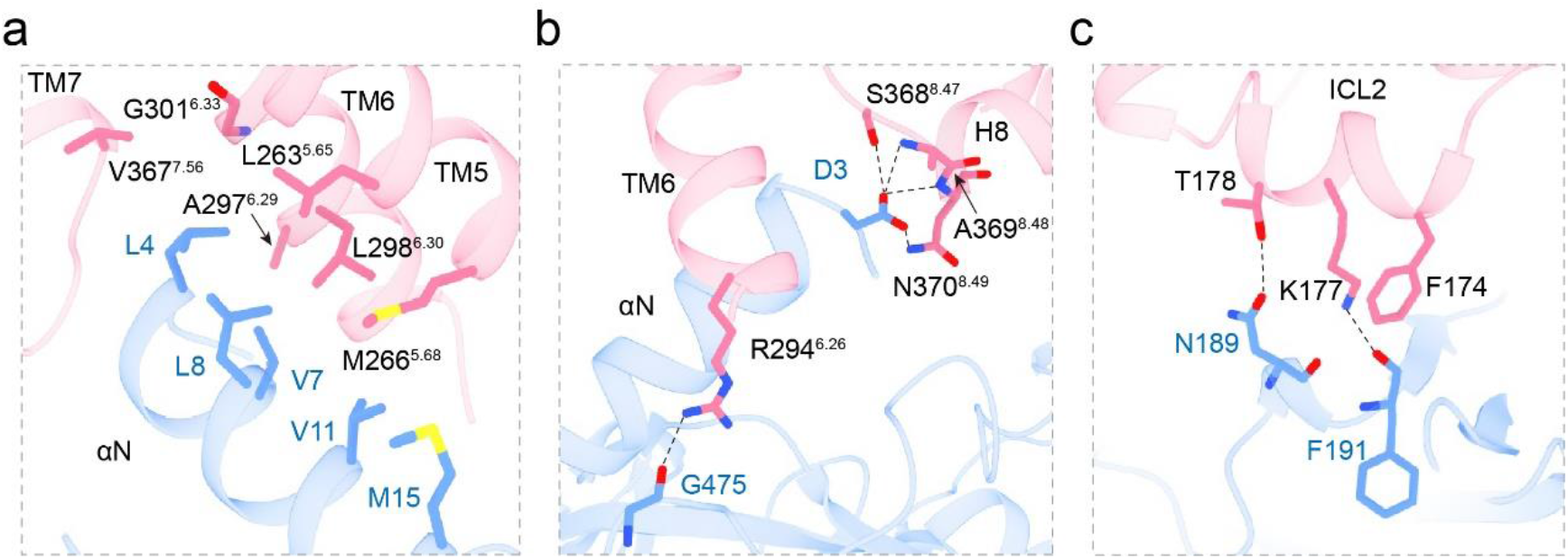
Interactions between NTSR1 and GRK2. **a, b,** Detailed interactions at the major interface between the NTSR1 cytoplasmic hydrophobic pocket and GRK2. **c**, Detailed interactions at the minor interface between the ICL2 from NTSR1 and GRK2. NTSR1 is shown in pink and GRK2 is shown in blue.

At the major interface, five hydrophobic residues (L4, V7, L8, V11, and M15) from the N-terminal helix of GRK2 form an extended hydrophobic patch, which is packed against a hydrophobic pocket formed by hydrophobic residues from TM5, TM6, and TM7 (L263^5.65^, M266^5.68^, A297^6.29^, L298^6.30^, G301^6.33^, and V367^7.56^) (Fig. 3a). In addition, the carboxylate side chain of D3 forms a network of hydrogen bonds with the main chain amine groups of A369^8.48^ and N370^8.49^, and the side chains of S368^8.47^ and N370^8.49^. R294^6.26^ also forms a direct hydrogen bond with the main chain carbonyl from G475 of GRK2 (Fig. 3b). These additional hydrogen bonds may also help to stabilize the N-terminal helix of GRK2 in the cytoplasmic pocket. At the ICL2 minor interface, F174 is packed against the main chain of N189 from GRK2, K177 forms a hydrogen bond with the main chain carbonyl of F191 from the GRK2 kinase domain, and T178 forms a hydrogen bond with the side chain of N189 (Fig. 3c). The total buried surface area between GRK2 and NTSR1 is 746 Å^2^, which is considerably smaller than the NTSR1-Gi interface of 1197 Å^2^, consistent with the relatively weak NTSR1-GRK2 interactions.

## The basis of SBI-553 biased agonism

SBI-553 is an arrestin-biased PAM ligand that specifically blocks G protein signaling but enhances arrestin signaling^7^. The high-quality density map clearly defines the binding mode of SBI-553 (Fig. 4), which adopts an inverted T-shape configuration and binds to the interface between NTSR1 and GRK2 (Fig. 4a, 4b). In the structure, SBI-553 forms extensive interactions with both receptor and GRK2 as summarized in Extended Data Table 2. Specifically, with the receptor, SBI-553 form predominately hydrophobic interactions with residues from TM2, TM3, TM5, TM6, TM7, and H8 (Fig. 4c, 4d). With GRK2, SBI-553 forms direct interactions with L4, E5, and L8 from the N-terminal helix (Fig. 4d), consistent with the enhanced binding of GRK2 to NTSR1 by SBI-553 (Fig. 1a, 1b).

**Fig. 4:**
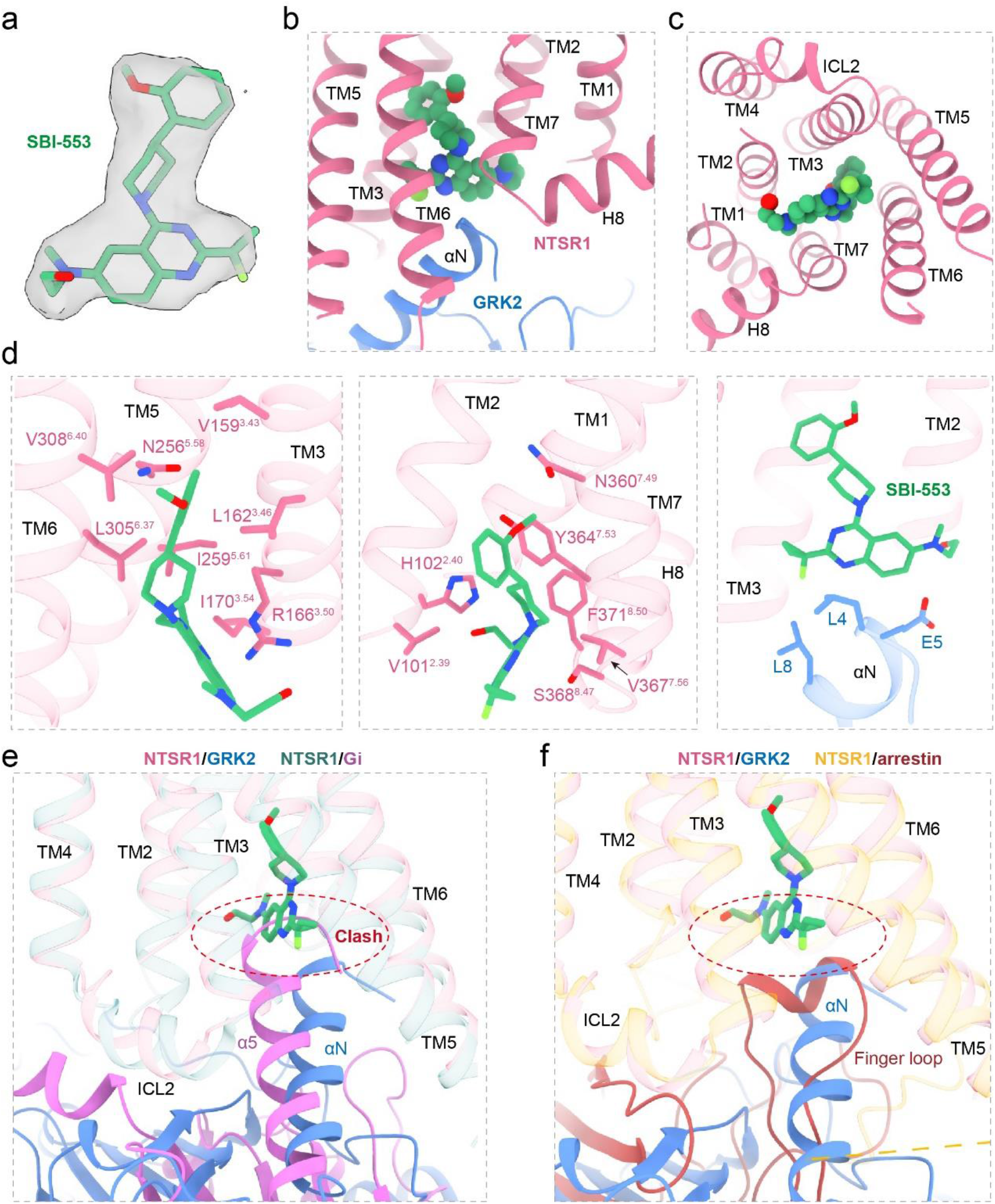
The binding mode of SBI-553 in NTSR1. **a,** The EM density of SBI-553, which is shown at a level of 0.12. **b, c,** The binding pocket of SBI-553 in NTSR1, from the front view (**b**) and top view (**c**). SBI-553 is highlighted in sphere. **d**, Detailed interactions between SBI-553 and NTSR1, as well as GRK2. SBI-553 is shown in green, NTSR1 is shown in pink, and GRK2 is shown in blue. **e,** Superposition of the NTSR1 from NTSR1-GRK2-Gαq complex and NTSR1-Gi complex showed the α5 of Gi protein would clash with SBI-553. **f,** Superposition of the NTSR1 from NTSR1-GRK2-Gαq complex and NTSR1-arrestin complex showed the finger loop of arrestin was compatible with the location of SBI-553.

The binding site of SBI-553 is unique and unexpected, which has not been observed in any GPCR structures determined to date^34,35^. Structure superposition of NTSR1 from its Gi complex onto the NTSR1-GRK2 structure reveal that the α5 helix from the G proteins occupies roughly the same space as occupied by the N-terminal helix of GRK2 (Extended Data Fig. 7). However, the α5 helix from the Gαi is up-shift by as much as 8.0 Å into the TMD pocket related to the N-terminal helix of GRK2 (Extended Data Fig. 7). In this orientation, the α5 helix from the G_αi_ would clash directly with the bound SBI-553 (Fig. 4e), thus providing a direct explanation for inhibition of G protein signaling by SBI-553. Importantly, structural superposition of NTSR1 from its arrestin complex reveals that the binding of SBI-553 would be compatible with arrestin binding to NTSR1 (Fig. 4f), consistent with its arrestin-biased signaling property.

## Universal features of GPCR-GRK interactions

In this paper, we have determined the structure of NTSR1 in complex with GRK2 and SBI-553 at a resolution of 2.92 Å, with the interface region approaching to 2.5 Å (Extended Data Fig. 2c, 2d). The relatively high resolution of the structure provides a clear binding mode of GRK2 and SBI-553 to NTSR1 as well as the mode of GRK2 membrane association. The primary binding site of GRK2 at NTSR1 is overlapped with the NTSR1 Gi binding site comprised by TM6, TM7 and H8, which structural features are highly similar in the active structures of various GPCRs (Extended Data Fig. 8), thus providing a basis for GRK2’s capability to interact with many different GPCRs.

In addition, the binding site of GRK2 at NTSR1 is the same as the GRK1 binding site in rhodopsin (Fig. 2c). In our structure, GRK2 has nearly complete structure with clear definition of many flexible regions, including the RHD and the active site tether (AST) loop, which tether the kinase domain in the active conformation. The residues from the N-terminal helix of GRK2 that interact with NTSR1 are highly conserved in all GRKs, suggesting the binding mode of GRK2 is a universal feature for all GRKs (Extended Data Fig. 6).

Finally, our structure reveals an unexpected binding mode of SBI-553, which is docked at the interface between GRK2 and NTSR1, consistent with its ability to enhance GRK2 binding to NTSR1 (Fig. 1a, 1b, 2b). The binding of SBI-553 is compatible with arrestin binding but would clash with G proteins (Fig. 4e, 4f), thus providing a direct mechanism for its arrestin-biased signaling capability. Together, our structure provides a solid model (Fig. 5) for understanding the details of GPCR-GRK interactions and biased signaling, and a basis for designing arrestin-biased ligands for NTSR1 and possibly other GPCRs.

**Fig. 5:**
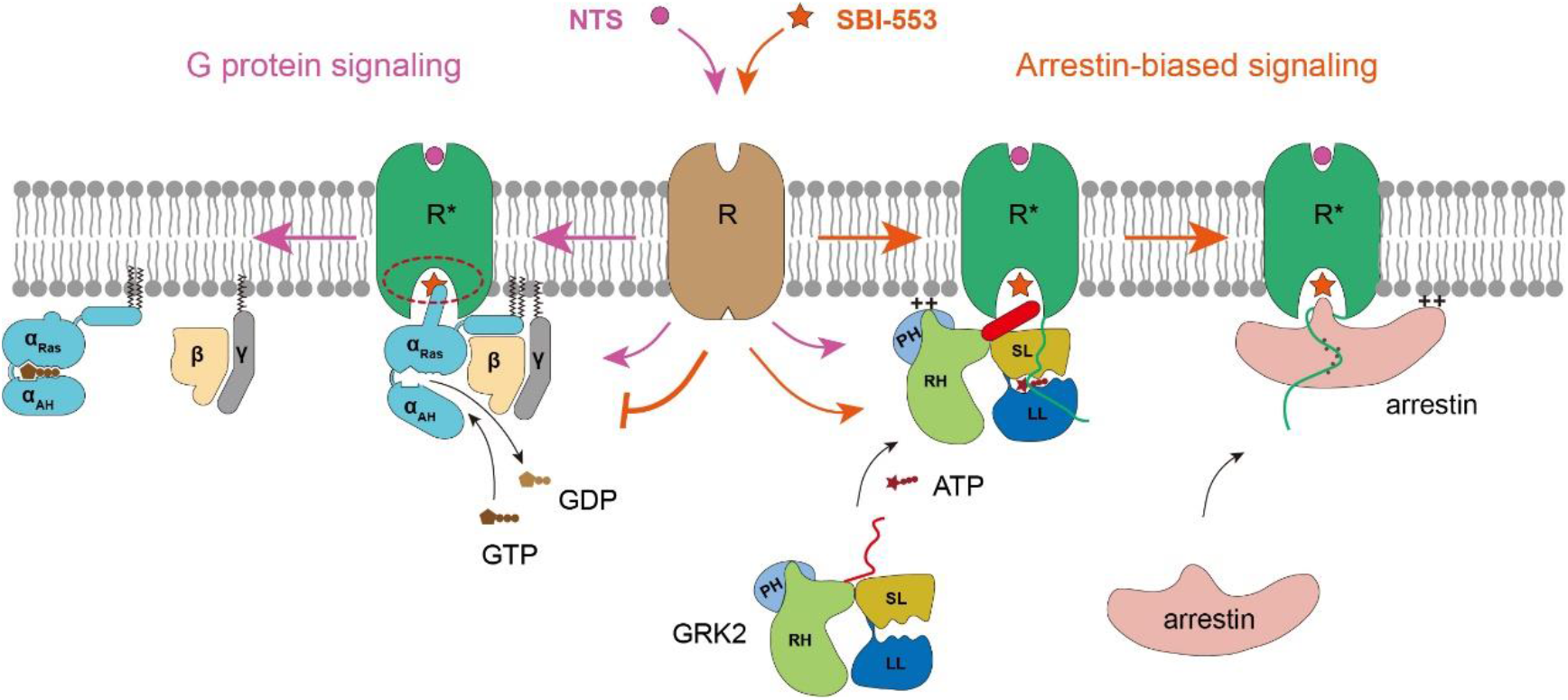
Cartoon presentation of NTSR1 signaling mediated by G protein and arrestin. NTS promotes NTSR1 to mediate both G protein and arrestin signaling but SBI-553 blocks G protein signaling and promotes GRK2 and arrestin signaling. ++ marks indicate membrane binding.

## Method

### Constructs

Human NTSR1 (residues 1-418) was codon-optimized for Sf9 expression and cloned into a modified pFastBac vector, which contains an N-terminal hemagglutinin (HA) signal peptide followed by a flag tag and a b562RIL (BRIL) epitope before the receptor. To improve the complex homogeneity and stability, the NanoBiT tethering strategy was applied by fusing a LgBiT subunit (Promega) at the receptor C-terminus after a GSSGGSGGGG linker^31,32^. Bovine GRK2 was cloned with a C-terminal GSSGGSGGGG linker followed by the HiBiT (peptide86) subunit^32^. Additionally, three mutations (A292P, R295I and S455D) were also incorporated into GRK2 by site-directed mutagenesis to enhance the affinity between GRK2 and Fab6^18^. Gαq construct was modified into a pFastBac vector. And the native N terminus (residues 1-28) of Gαq was replaced with Gαi1 to facilitate the expression of Gαq^16^.

### Expression and purification of NTSR1-GRK2-Gαq complex

NTSR1-LgBiT, Gαq, GRK2-HiBiT and Ric8a (a gift from Brian Kobilka) were coexpressed in *Sf9* insect cells (Invitrogen) using the Bac-to-Bac baculovirus expression system (ThermoFisher). Cell pellets were thawed and lysed in 20 mM HEPEs, pH 7.4, 100 mM NaCl, 10% glycerol, 10 mM MgCl_2_, 10 mM NaF and 30 μM AlCl3 supplemented with Protease Inhibitor Cocktail, EDTA-Free (TargetMol). The NTSR1-GRK2-Gαq complex was formed in membranes by the addition of 10 μM NTS (Genscript), 10 μM saturosporine, 10 μM SBI-553 (TargetMol) and 50 μM GDP. The suspension was incubated for 0.5 h at room temperature before centrifugation at 80,000 × g for 30 min. The membrane was then resuspended with the same buffer and solubilized using 0.5% (w/v) n-dodecyl β-D-maltoside (DDM, Anatrace), 0.1% (w/v) cholesterol hemisuccinate (CHS, Anatrace) for 2 h at 4 °C. The supernatant was collected by centrifugation at 80,000 × g for 40 min and then incubated with G1 anti-Flag affinity resin (Genscript) for 2 h at 4 °C. After batch binding, the resin was loaded into a plastic gravity flow column and washed with 20 column volumes of 20 mM HEPEs, pH 7.4, 100 mM NaCl, 10% glycerol, 10 mM MgCl_2_, 10 mM NaF, 30 μM AlCl_3_, 10 μM NTS, 10 μM saturosporine, 10 μM SBI-553 and 50 μM GDP, 0.01% (w/v) DDM, 0.002%(w/v) CHS, and 0.05%(w/v) digitonin, further eluted with 10 column volumes of the same buffer plus 0.2 mg/mL Flag peptide. The complex was then concentrated using an Amicon Ultra Centrifugal Filter (MWCO 100 kDa) and injected onto a Superose 6 Increase 10/300 GL column (GE Healthcare) equilibrated in the buffer containing 20 mM HEPEs, pH 7.4, 100 mM NaCl, 10 mM MgCl_2_, 10 mM NaF, 30 μM AlCl3, 10 μM NTS, 10 μM saturosporine, 5 μM SBI-553, 50 μM GDP, and 0.03% (w/v) digitonin. To stabilize the NTSR1-GRK2-Gαq complex, the peak fractions were collected and crosslinked using 0.01 mM BS_3_ for 0.5 h at room temperature, stopped crosslinking by addition of 80 mM of glycine, and then concentrated to approximately 10 mg/ml for cryo EM analysis.

### Cryo-EM grid preparation and data collection

For the preparation of cryo-EM grids, 3 μL of the purified protein at 10 mg/mL were applied onto a glow-discharged holey carbon grid (CryoMatrix Amorphous alloy film R1.2/1.3, 300 mesh). Grids were plunge-frozen in liquid ethane using Vitrobot Mark IV (Thermo Fischer Scientific). Frozen grids were transferred to liquid nitrogen and stored for data acquisition. Cryo-EM imaging of the complex was performed on a Titan Krios at 300 kV in the Advanced Center for Electron Microscopy at Shanghai Institute of Materia Medica, Chinese Academy of Sciences (Shanghai China).

A total of 57,477 movies for the NTSR1-GRK2-Gαq complex were collected by a Gatan K3 Summit direct electron detector with a Gatan energy filter (operated with a slit width of 20 eV) (GIF) at a pixel size of 0.824 Å using the EPU software. The micrographs were recorded in counting mode with a defocus ranging from −1.2 to −2.2 μm. The total exposure time was 3.33 s with a dose of 50 electrons, and intermediate frames were recorded in 0.104 s intervals, resulting in a total of 36 frames per micrograph.

### Image processing and map construction

A total of 57,477 dose-fractioned movies were used for correction of beam-induced movement using a dose-weighting scheme in MotionCor2^36^ and their contrast transfer function parameters were estimated by Patch CTF estimation in CryoSPARC^37^. For the NTSR1-GRK2-Gαq complex, particle selection was performed by blob picking using CryoSPARC^37^, and 40,940,867 particles were extracted and further subjected to an initial reference-free 2D classification. Interactive 2D and 3D classifications were performed to discard poorly defined particles, and 1,149,932 particles were retained. These particles were divided into 6 subclasses using Ab-initial model and heterorefinement, resulting in two subsets with complete NTSR1-GRK2-Gαq complex. Two maps from the two subsets showed slightly difference especially relative position of GRK2 and Gαq. We merged the two subsets, and performed another round of Ab-initial model and hetero-refinement to remove particles without clear NTSR1-GRK2-Gαq complex. In the 5 subclasses, two well-defined subsets, containing 287,853 particles and 216,282 particles, respectively, were subsequently subjected to non-uniform refinement in CryoSPARC^37^, generated two different maps with global resolution of 2.92 Å and 3.09 Å. Resolution was estimated in the presence of a soft solvent mask and based on the gold standard Fourier shell correlation (FSC) 0.143 criterion. Local resolution was estimated in cryoSPARC^37^ using default parameters. Unless indicated otherwise, the maps shown in figures were sharpened with B factors estimated in the nonuniform refinement.

To analyze the flexibility of the NTSR1-GRK2-Gαq complex, we performed cryoSPARC 3D variability analysis (3DVA)^38^. The 3DVA was performed with mask on the complex, generated from non-uniform refinement. The 3DVA was analyzed across three principal components that estimated the most common motions. One of the components showed pronounced motion between GRK2 and Gαq and the movie that consisted of 20 volume frame data were presented by Chimera (v1.4) in the Extended Data movie.

### Model building and refinement

For the NTSR1-GRK2-Gαq complexes, the AlphaFold model of NTSR1 and the structure of G Protein-Coupled Receptor Kinase 2 in Complex with Gαq and Gβγ Subunits (PDB code: 2BCJ), were used as the start for model rebuilding and refinement against the electron microscopy map. The model was docked into the EM density map using Chimera^39^, followed by iterative manual adjustment and rebuilding in COOT^40^ and ISOLDE^41^. Real space and reciprocal space refinements were performed using Phenix^42^ programs with secondary structure and geometry restraints. The final refinement statistics were validated using the module “comprehensive validation (cryo-EM)” in Phenix^42^. The final refinement statistics are provided in Extended Data Table 1. Structure figures were prepared in ChimeraX^43^ and PyMOL (https://pymol.org/2/).

### Calculation of NTSR1-Gi protein and NTSR1-GRK2 interface area

NTSR1-GRK2-Gαq complex and NTSR1-Gi protein complex (PDB ID: 6OS9) were used for the calculation of NTSR1-GRK2, NTSR1-Gi interface areas respectively, using PDBePISA web server (<u>PDBe < PISA < EMBL-EBI</u>). During the process, NTSR1-GRK2-Gαq complex and NTSR1-G protein complex were uploaded, and the Accessible surface area (ASA) calculations are based on finite element analysis through the “interface” module.

### NanoBiT assay for GRK2 recruitment

The full-length NTSR1 (1–418) was cloned into pBiT1.1 vector (Invitrogen) with a FLAG tag at its N-terminus and LgBiT at its C-terminus. Bovine GRK2 (residues 1-689) was cloned into pBiT2.1 vector (Invitrogen) with a modified SmBiT (peptide104: MVEGYRLFEKIS)^31^ and a GSSGGGGSGGGGSSG linker at its N-terminus. AD293 cells were cultured in DMEM/high Glucose (GE healthcare) supplemented with 10% (w/v) FBS (Gemini). Cells were maintained at 37 °C in a 5% CO_2_ incubator with 300,000 cells per well in a 6-well plate. Cells were grown overnight and then transfected with 1.5 μg NTSR1 and 1.5 μg GRK2 constructs by FuGENE^®^ HD transfection reagent in each well for 24 h. Cells were harvested and re-suspended in Hanks’ balanced salt solution buffer (HBSS) at a density of 5 × 10^5^ cells/ml. The cell suspension was seeded in a 384-well plate at a volume of 10 μl per well, followed by 10 μl HBSS or 10 μl HBSS containing 1 μM SBI-553, 10 μl HBSS containing different concentrations of NTS, and another 10 μl the NanoLuc substrate (furimazine, 1:25 dilution, Promega) diluted in the detection buffer. The luminescence signal was measured with an EnVision plate reader at room temperature.

### Tango assay

Human NTSR1 (1-418) was cloned into pcDNA6 vector consisting of an expression cassette with tobacco etch virus (TEV) protease cleavage site and the transcriptional activator tTA at the C terminus. A TEV protease cDNA was fused to the C-terminus of GRK2 (1-689). Interaction between NTSR1 and GRK2 leads to the cleavage of the TEV site, thus releasing tTA to trigger tTA-dependent luciferase reporter gene expression. For Tango assays, HTL cells were cultured in 24-well plate at a density of 5 ×10^4^ cells/well for 24 h, and then transfected with 10 ng NTSR1, 10 ng GRK2 plasmids and 5 ng of phRG-tk Renilla luciferase expression plasmids using FuGENE^®^ HD transfection reagent. After transfection for 24 h, cells were incubated overnight with PBS (vehicle), or different concentrations of ligands. Then luciferase activities were evaluated according to manufacturer’s protocols of the Dual Luciferase Kit (Promega).

## Acknowledgments

The cryo-EM data were collected at the Advanced Center for Electron Microscopy, Shanghai Institute of Materia Medica (SIMM). The authors thank the staff at the Advanced Center for Electron Microscopy for their technical support. This work was partially supported by Ministry of Science and Technology (China) grants (2018YFA0507002 to H.E.X.); Shanghai Municipal Science and Technology Major Project (2019SHZDZX02 to H.E.X.); Shanghai Municipal Science and Technology Major Project (H.E.X.); CAS Strategic Priority Research Program (XDB37030103 to H.E.X.); the National Natural Science Foundation of China (32130022 to H.E.X., 32171187 to Y.J., 82121005 to H.E.X. and Y.J.).

## Author contributions

J.D. designed the expression constructs, purified the proteins, performed cryo-EM grid preparation and data collection, participated in functional studies, participated in figure and manuscript preparation; H.L. performed cryo-EM data calculations, model building, and participated in figure preparation; Y-J.J participated in protein purification and functional studies; Q.Y. and K.W. participated in cryo-EM data calculations, X.L., W.Y., S.Z., and T.G. participated in the experiments; Y.J. supervised the studies, and participated in manuscript preparation; H.E.X. and J.D. conceived the project, analyzed the structures, and wrote the manuscript with inputs from all authors.

---

**Extended Data Fig. 1.**
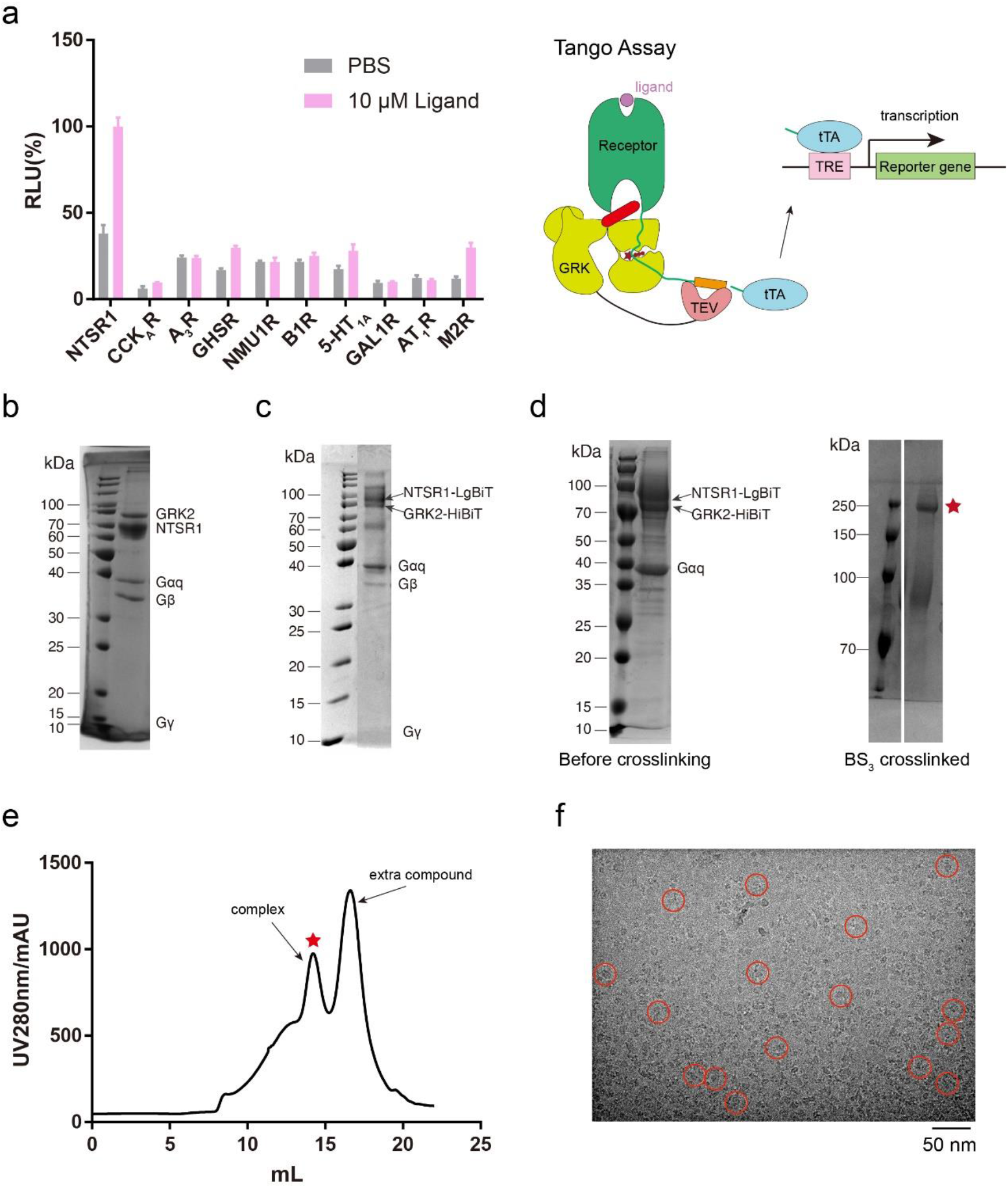
NTSR1-GRK2-Gαq complex assembly. **a**, Screening for GPCR– GRK2 complexes by tango assay. RLU, relative luciferase units, which was normalized to the values of NTSR1. **b-d** SDS-PAGE of the complexes. NTSR1-GRK2-Gαq-Gβγ complex (**b**), NTSR1_LgBiT-GRK2_HiBiT-Gαq-Gβγ complex (**c**), NTSR1-GRK2-Gαq complex before crosslinking (left panel of **d**) and NTSR1-GRK2-Gαq complex crosslinked by BS_3_ (right panel of **d**). **e,** Size-exclusion chromatography elution profile of the NTSR1-GRK2-Gαq complex. Red star indicates the monomer peak of the complex. **f,** Cryo-EM micrograph of the NTSR1-GRK2-Gαq complex. Particles picked for 3D classifications were highlighted in red circles.

**Extended Data Fig. 2.**
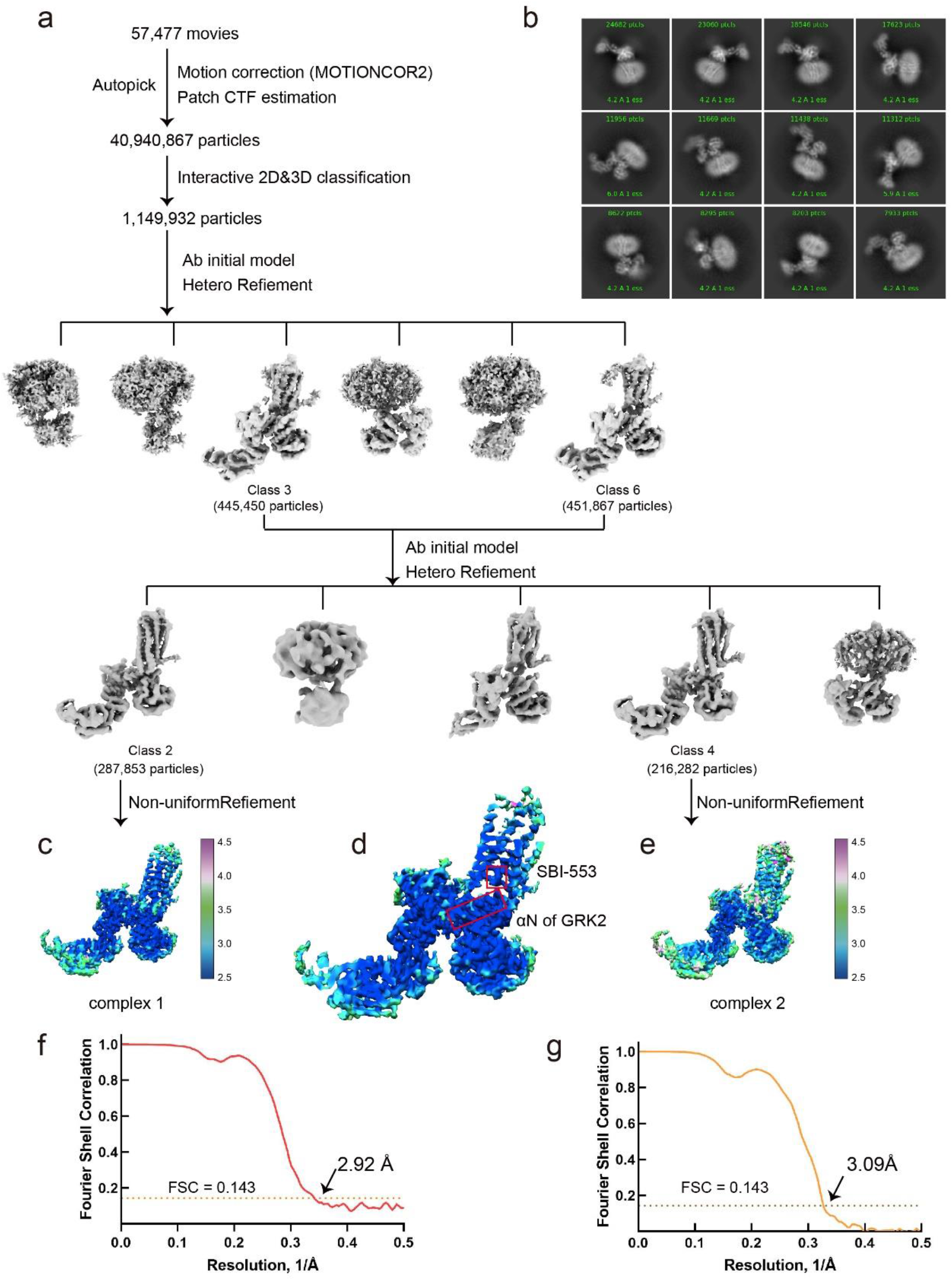
Single-particle reconstruction of the NTSR1-GRK2-Gαq complex. **a**, Flowchart of cryo-EM data analysis of the NTSR1-GRK2-Gαq complex. **b,** Micrograph of the reference-free 2D class averages. **c-e,** Two cryo-EM maps of the NTSR1-GRK2-Gαq complexes were generated and colored by local resolutions from 2.5 Å (blue) to 4.5 Å (purple) (**c and e**). The first map is enlarged for the clarity of SBI-553 and αN of GRK2. SBI-553 and αN of GRK2 are highlighted in red squares (**d**). **f, g,** The “Gold-standard” Fourier shell correlation (FSC) curve indicates that the overall resolution of the electron density map of the NTSR1-GRK2-Gαq complex 1 is 2.92 Å, the NTSR1-GRK2-Gαq complex 2 is 3.09 Å.

**Extended Data Fig. 3.**
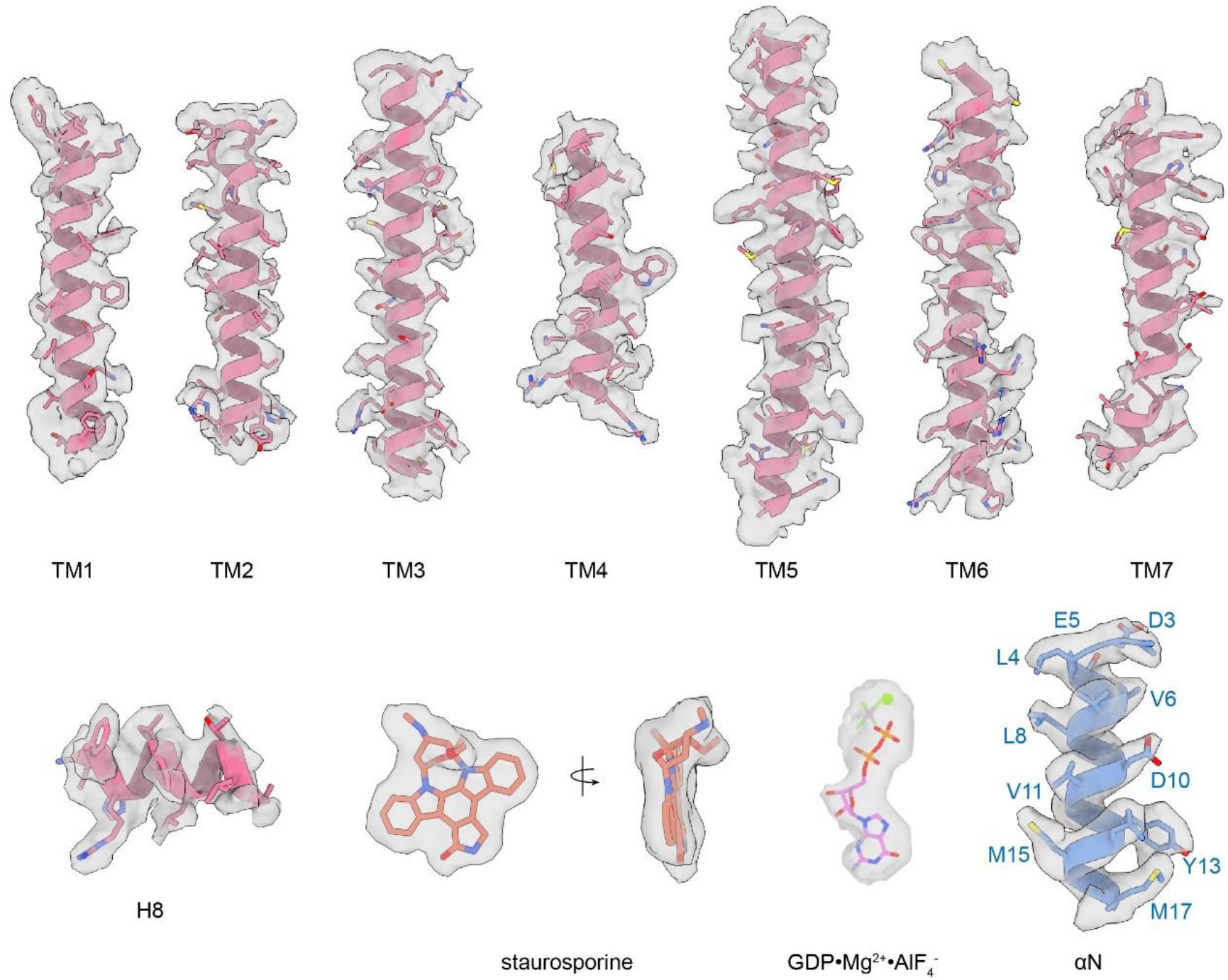
Cryo-EM density maps with all transmembrane helices, and H8 of NTSR1, staurosporine, GDP·Mg^2+^·AlF4^-^ and αN of GRK2.

**Extended Data Fig. 4.**
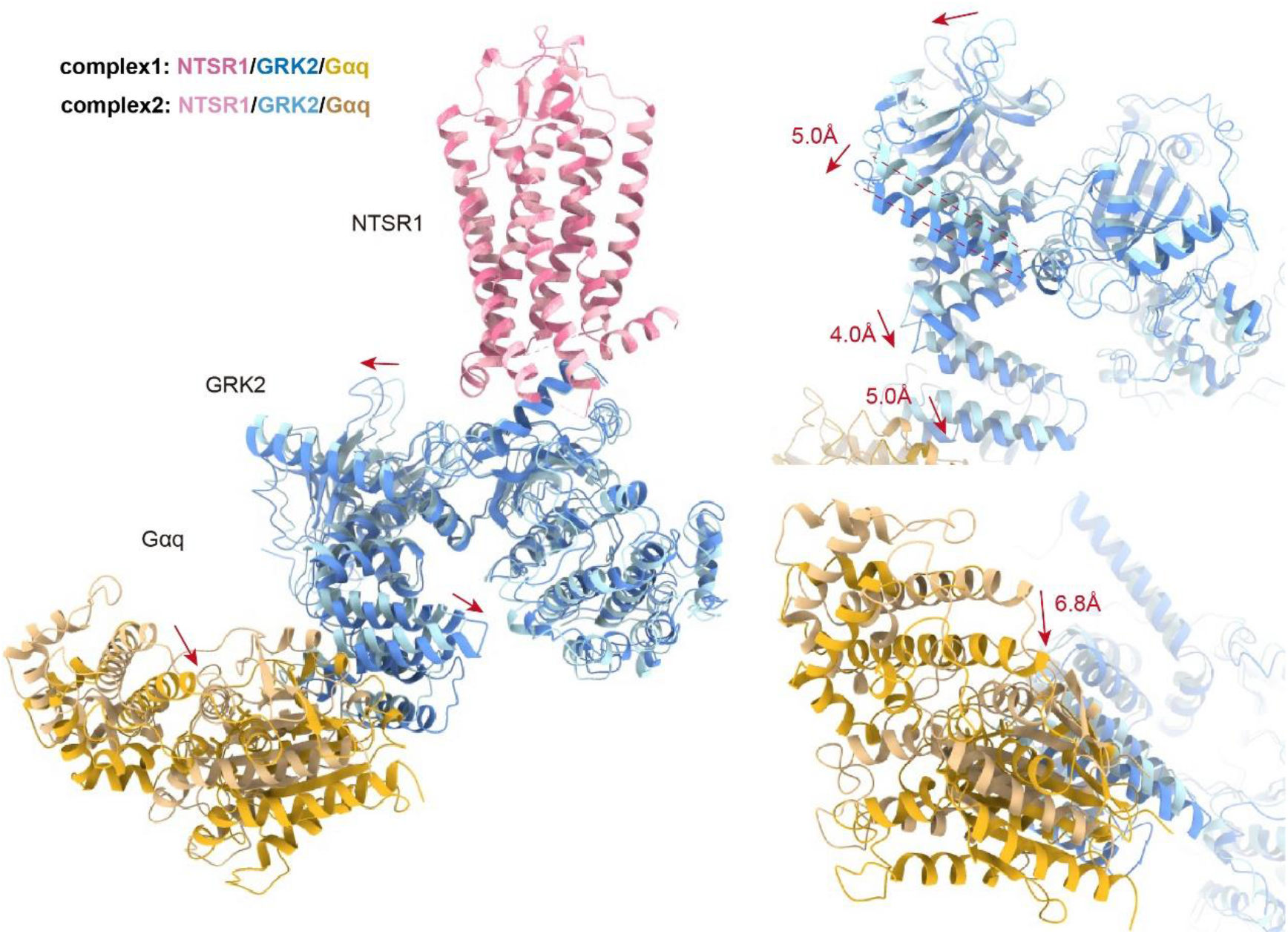
Structural comparison of the NTSR1-GRK2-Gαq complex1 and 2. Comparison of these two complexes reveals that they have very similar NTSR1 structure but a swing of GRK2 and Gαq of ~5-6 Å related to NTSR1.

**Extended Data Fig. 5.**
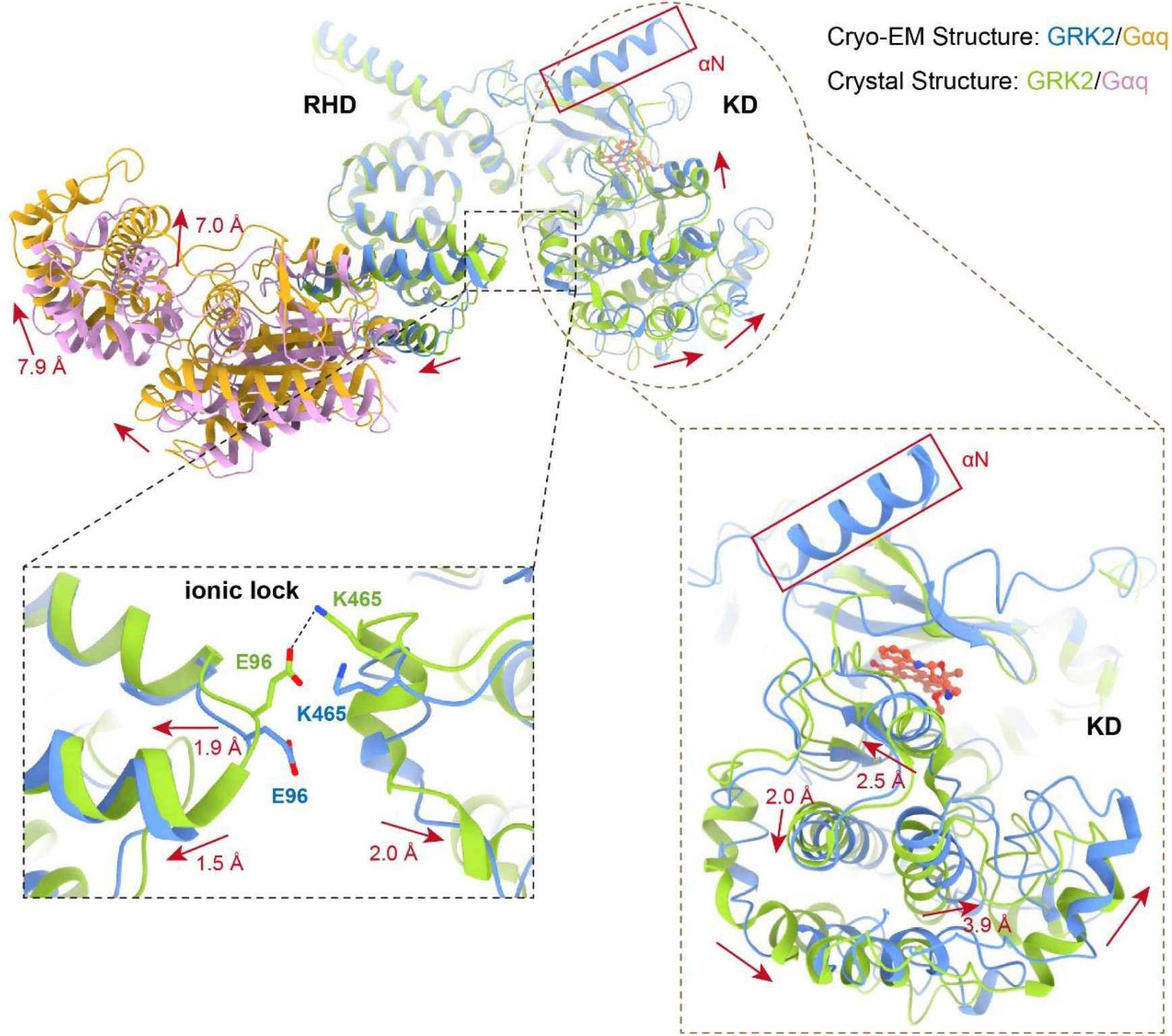
Structural comparison of the GRK2-Gαq from the cryo-EM structure NTSR1-GRK2-Gαq complex with the crystal structure of GRK2-Gαq-Gβγ. Comparing the GRK2 structure from the NTSR1 complex to the crystal structure of GRK2 from the complex with Gαq and Gβγ reveals three major differences. The GRK2 structure from the NTSR1 complex contains a N-terminal helix that is packed onto the kinase domain, has a breakage in the ionic lock between its RHD from the KD, and adopts a closed conformation in its KD by 2-3 Å shifts of the KD relative to the KD of GRK2 from the crystal structure.

**Extended Data Fig. 6.**
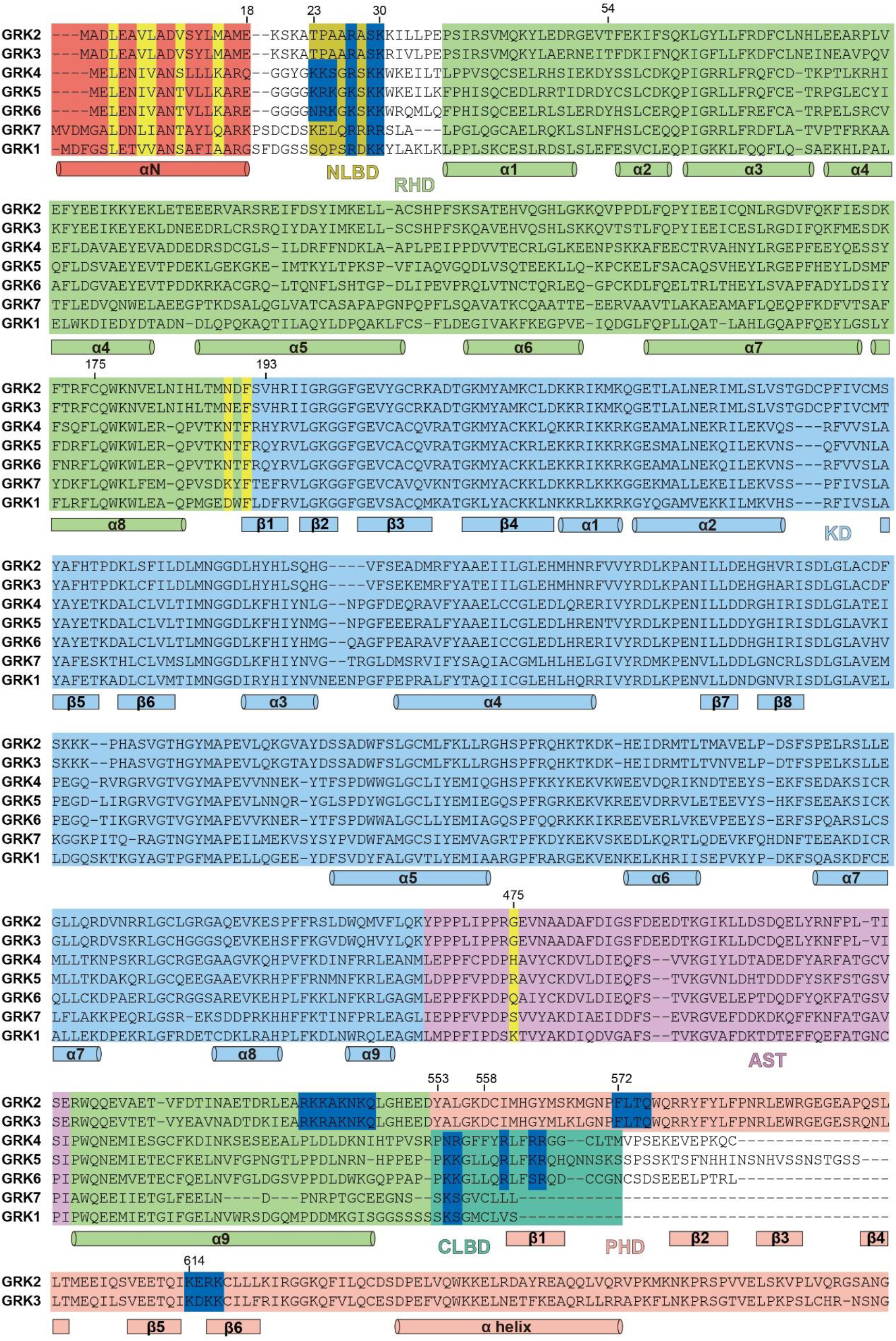
Sequence alignment of human GRKs. The N-terminal helix (αN) is highlighted in red. The RHD is highlighted in green and KD is in light blue. The AST loop extended from the kinase domain is in light purple. The PHD of GRK2 and GRK3 are in pink. N-terminal lipid binding domain (NLBD) and C-terminal lipid binding domain (NLBD) according to GRK5 are highlighted in dark yellow and light green, respectively. Residues that may interact with membrane lipid are highlighted in dark blue. And residues from GRK2 that interact with NTSR1 are highlighted in yellow, α, α helix. β, β strand.

**Extended Data Fig. 7.**
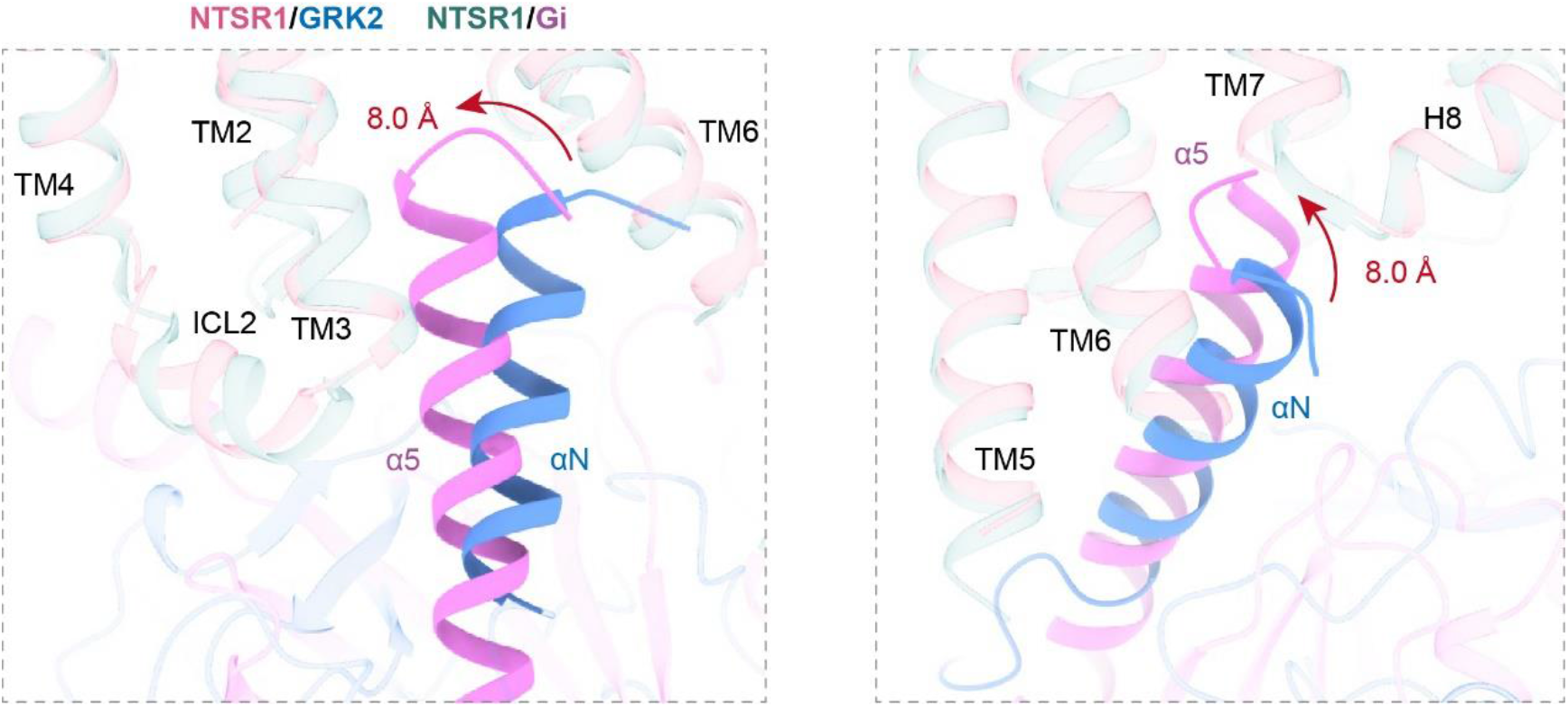
Structural comparison of NTSR1-GRK2-Gαq complex with NTSR-Gi complex. Superposition of the NTSR1 from NTSR1-GRK2-Gαq complex and NTSR1-Gi complex showed the α5 helix from the Gαi is up-shift by 8.0 Å into the TMD pocket related to the N-terminal helix of GRK2.

**Extended Data Fig. 8.**
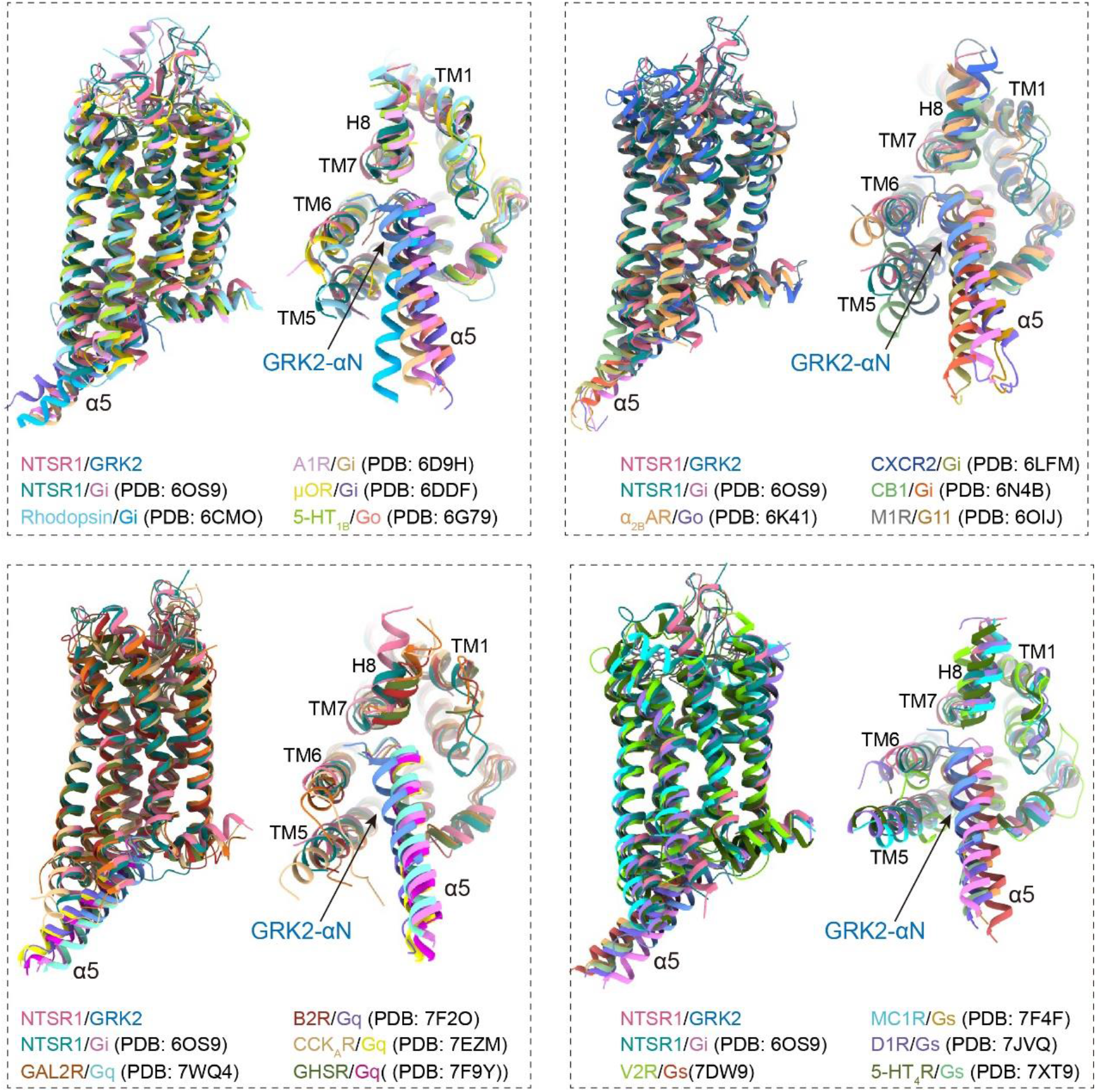
Structural comparison of NTSR1-GRK2-Gαq complex with NTSR-Gi complex and other GPCR-G protein complexes. Superposition of the receptors from different GPCR complexes showed that the active GPCRs had very similar 3D architecture, and the location of α5 helix from different G proteins overlapped with the N-terminal helix of GRK2.

**Extended Data Table 1.**
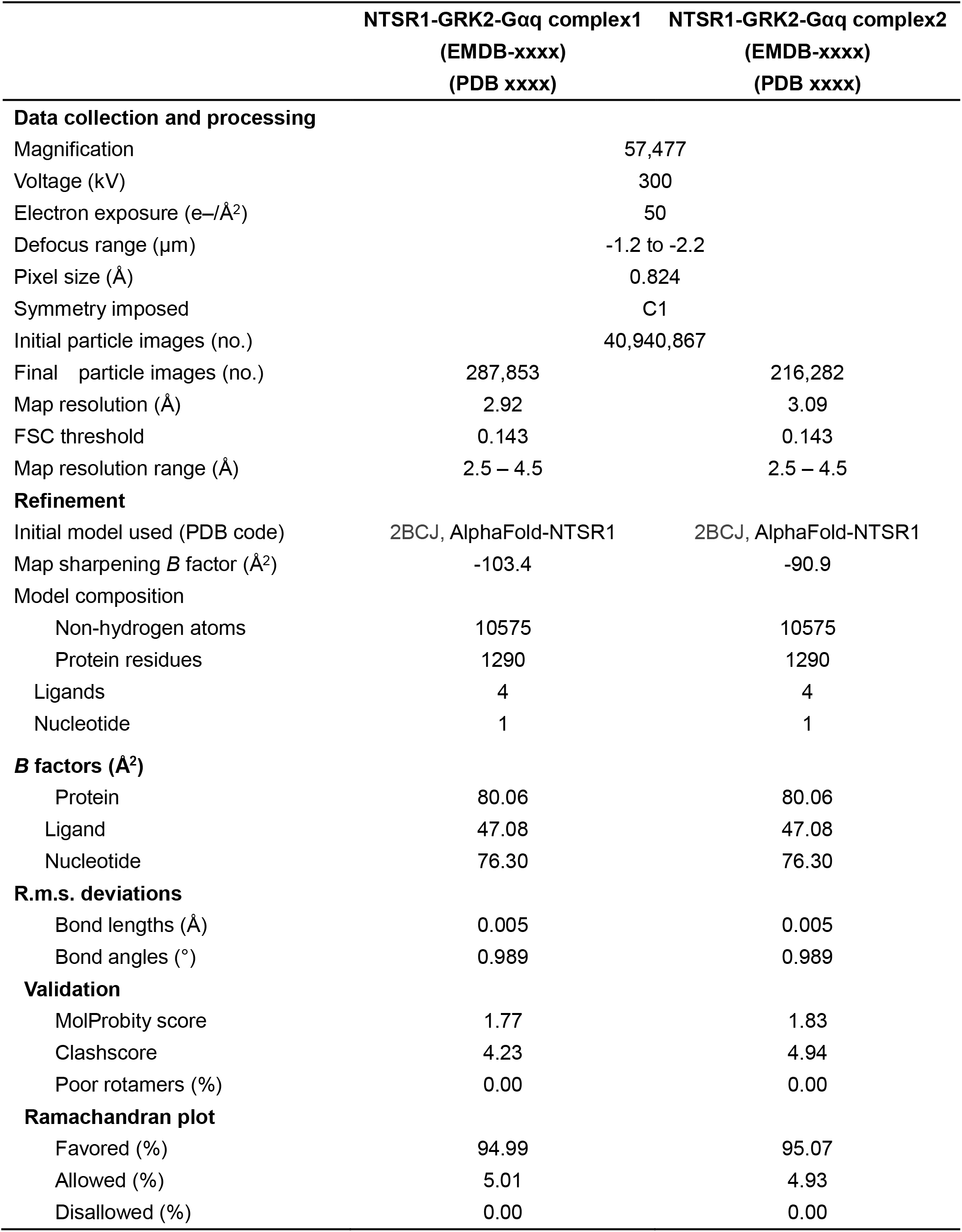
Cryo-EM data collection, refinement and validation statistics.

**Extended Data Table 2.** Interactions of SBI-553 with NTSR1 and GRK2.

|  |  |  |
| --- | --- | --- |
| SBI-553 | V101<sup>2.39</sup> | NTSR1 |
|  | H102<sup>2.40</sup> |  |
|  | V159<sup>3.43</sup> |  |
|  | L162<sup>3.46</sup> |  |
|  | R166<sup>3.50</sup> |  |
|  | I170<sup>3.54</sup> |  |
|  | N256<sup>5.58</sup> |  |
|  | I259<sup>5.61</sup> |  |
|  | L305<sup>6.37</sup> |  |
|  | V308<sup>6.40</sup> |  |
|  | N360<sup>7.49</sup> |  |
|  | Y364<sup>7.53</sup> |  |
|  | V367<sup>7.56</sup> |  |
|  | S368<sup>8.47</sup> |  |
|  | F371<sup>8.50</sup> |  |
|  | L4 | GRK2 |
|  | E5 |  |
|  | L8 |  |

